# Supervised Promoter Recognition: A benchmark framework

**DOI:** 10.1101/2021.09.07.459345

**Authors:** Raul Ivan Perez Martell, Alison Ziesel, Hosna Jabbari, Ulrike Stege

## Abstract

Deep learning has become a prevalent method in identifying genomic regulatory sequences such as promoters. In a number of recent papers, the performance of deep learning models have continually been reported as an improvement over alternatives for sequence-based promoter recognition. However, the performance improvements in these models do not account for the different datasets that models are being evaluated on. The lack of a consensus dataset and procedure for benchmarking purposes has made the comparison of each model’s true performance difficult to assess.

We present a framework called Supervised Promoter Recognition Framework (‘SUPR REF’) capable of streamlining the complete process of training, validating, testing, and comparing promoter recognition models in a systematic manner. *SUPR REF* includes the creation of biologically relevant benchmark datasets to be used in the evaluation process of deep learning promoter recognition models. We showcase this framework by comparing the models’ performance on alternative datasets, and properly evaluate previously published models on new benchmark datasets. Our results show that the reliability of deep learning *ab initio* promoter recognition models on eukaryotic genomic sequences is still not at a sufficient level, as precision is severely lacking. Furthermore, given the observational nature of these data, cross-validation results from small datasets need to be interpreted with caution.

**Availability:** Source code and documentation of the framework is available online at https://github.com/ivanpmartell/suprref

## 1 Introduction

Promoters are non-coding genomic elements necessary for the expression of their associated gene(s). Most promoters include transcription factor binding sites (TFBS): short, often palindromic DNA sequences bound by transcription factor proteins to provide greater control over a promoter’s activity. TFBS can be summarised as ‘motifs’ (Lambert et al. [2018]), that represent the set of related short sequences preferred by a given Transcription Factor (TF). An example of a motif found in eukaryotic promoters is the TATA box, a cis-regulatory element characterised by its consensus sequence of repeating T and A base pairs. Other relevant promoter motifs are in the JASPAR database (Fornes et al. [2019]). A minimal promoter must be able to recruit RNA polymerase (RNAP) to allow for transcription to occur. Promoters can be separated into three main regions: core, proximal, and distal. Core promoters are located closest to their gene, containing the transcription start site (TSS) of the gene and usually include general TFBS to create an RNAP binding site. Proximal promoters are located approximately 200 base pairs upstream from the TSS, and are the DNA regions where more gene-specific TF bind. Distal promoters are generally found thousands of base pairs upstream from the core promoter and include several other TFBS that can recruit proteins to enhance or silence the RNAP’s transcription process. While a promoter sequence itself is not expressed, mutations to the promoter sequence can have prominent impact on gene expression. E.g., recombination events may occur, which produce a novel promoter-gene combination, resulting in a significantly different expression pattern for the controlled gene; these are observed in certain types of cancer (Krzyzanowski et al. [2019]). As the central regulatory feature for gene expression, promoters can provide potentially significant information used for predicting downstream gene expression patterns. Rudge et al. [2016] have shown that understanding gene expression patterns can lead to therapies for disease control or prevention. Therefore, understanding and reliably identifying promoters within genomes are essential to this goal.

Identification of promoters through their genomic sequence is highly complex because of their sequence and structure diversity, which excludes any universal promoter elements. Considering the central dogma of molecular biology, all of the non-genetic effects involved in promoter activity could, in theory, be ultimately mapped to the DNA. This allows the genetic sequence of promoters, transcription factors, and other interacting proteins available in the genome to have the ability to fully characterise a promoter. Therefore, we focus on the promoter identification problem known as *ab initio promoter recognition*, which entails the use of DNA sequences solely to identify promoters. The work presented here can aid in any type of ab initio promoter recognition, but initially aims to help in recognition models for eukaryotic RNAP II promoters since they transcribe RNAs that will become messenger RNAs (mRNAs) and also small regulatory RNAs — the former being the products used by ribosomes to synthesize proteins, while the latter play a role in regulatory processes such as activation and inhibition of gene transcription.

A review by Li et al. [2015] reveals that current promoter identification efforts mainly lie in supervised machine learning (ML) techniques, with deep learning (DL) as the latest promising approach by utilizing data from different High-Throughput Sequencing (HTS) methods. Early ML models require the use of biological signals to be ‘handcrafted’ into features. Crafting these features can become especially problematic when domain-knowledge is incomplete, as is the case in promoter recognition. DL can account for incomplete domain-knowledge by ‘learning’ these features, while additional ‘handcrafted’ features can be supplied when domain-knowledge is present. Once *ab initio* methods can accurately recognise promoters, the methods and their outputs can be analysed to understand the sequences affecting specific promoters, their relation to genetic activity, and provide further insights into the genomic characteristics of promoters.

Performance of all ML-based methods is heavily impacted by the underlying training data used. Raeder et al. [2012] demonstrate that this problem can be exacerbated when using imbalanced datasets, such as promoter recognition. This imbalance is evident, as promoter sequences generally account for less than 1% of a genome. Since we are interested in the minority class (promoters) which contains significantly fewer instances, models will tend to focus on learning the characteristics of the majority class (non-promoters), therefore neglecting to learn promoter features. Scientists must ensure that the training data used in the ML process is a statistically representative sample of the many genomic sequences found in nature. It is also crucial that a held-out dataset be used for testing and comparing different models’ performance. This means that a trained model must not have previously seen any data within the testing dataset to ensure the validity of its performance metrics. Results from common evaluation methods for an ML model, such as cross-evaluation, can be especially deceptive when the overall dataset used for training and testing is limited. Therefore, results obtained in this manner can create the impression that a model’s performance is adequate when in reality this is not the case. Early ML models for *ab initio* promoter recognition were benchmarked by Bajic et al. [2006]. The benchmarking process was done using a limited subset of available genomic data, and found the models to be insufficiently sensitive for promoter recognition. Deep learning promoter recognition (DLPR) models frequently exhibit improved recognition performance over previous models through cross-validation assessments. The caveat is that these models are still being trained and tested on similarly limited datasets, generally comprising less than 60,000 DNA sequences, which cannot possibly cover a representative sample of the many genomic sequences found in nature.

We present a framework called *Supervised Promoter Recognition Framework* (‘SUPR REF’) capable of streamlining the complete process of training, validating, testing, and comparing promoter recognition models in a systematic manner. Our framework includes two different but compatible methods for the creation of benchmark datasets, to be used in the evaluation process of DLPR models. The first method utilises sequence alignment to find the promoter sequences in a genome, while the second uses promoter annotations that specify the TSS to extrapolate a promoter’s location within the genome. In addition, we implemented previously published models where no source code was provided to help ease reproducible research within the field. We benchmark published DL promoter recognition models using *SUPR REF* and assess their performance on realistic genome datasets.

We demonstrate the benchmarking procedure of *SUPR REF* on three previously published studies: CNNProm by Umarov and Solovyev [2017], ICNNP by Qian et al. [2018], DProm by Oubounyt et al. [2019]. We also include more thorough experimentation on the most advanced architecture between them (DProm)—a convolutional recurrent neural network. Benchmarking of previous work includes a comparison between their training and testing datasets in addition to comparisons of strategies to create synthetic datasets. Experiments include multiple training and testing datasets showcasing a comparison of different promoter annotations, sampling methods to overcome the imbalanced data, and the effect of output functions in neural networks.

We show that the reliability of deep learning *ab initio* promoter recognition models on eukaryotic genomic sequences is still not at an acceptable level, as precision is severely lacking. We do this by utilising *SUPR REF* to create larger biologically relevant datasets to test models more thoroughly. The remainder of the paper is organized as follows. Section 2 describes *SUPR REF* and the benchmarks performed to obtain our results. In Section 3, we examine the results from our implementations and their comparisons, along with the benchmarking results. Furthermore, we provide results of experiments on interesting functionality within DLPR models and analysis of datasets that provide these models with biased performance metrics. Finally, Section 4 recapitulates our research aims and its significance, as well as suggesting approaches for future work.

## 2 Materials and methods

We introduce our framework *SUPR REF* and the implemented models and datasets from previously published studies to showcase its use. We then introduce larger benchmarking datasets that reflect the statistics of promoter sequences found in nature, and suggest a method for a fair benchmarking of the DLPR models. Finally, we describe the metrics within the evaluation procedure for model comparison and benchmarking.

### 2.1 SUPR REF

Different frameworks exist for generalised use of DL in bioinformatics (Chen et al. [2019], Kopp et al. [2020]), however their use in more specific tasks would require extensive re-engineering. While some specialised frameworks exist in other domains (see Budach and Marsico [2018], Avsec et al. [2019]), to the best of our knowledge none exists for promoter recognition. A promoter recognition framework should aid in rapid advancement of novel recognition techniques, but must also offer benchmarking capabilities that can clearly assess the significance of these advances over previous models. It is important for the benchmarking process to be simple and systematic to limit the amount of time spent during this process and to help researchers focus on the creation of better models. These capabilities can be achieved using *SUPR REF*.

*SUPR REF* offers an advantage of domain-knowledge datasets and models that can account for promoter-specific problems that would otherwise go unnoticed in a generalised setting. Additionally, *SUPR REF* allows researchers without prior DL knowledge to compare models in a systematic way and to make informed decisions of which model to use within their specific context.

*SUPR REF* is a Python 3 framework designed to ease the training and testing procedures of machine learning models for promoter recognition. The framework can be installed as a command-line tool for automation and productivity purposes. It contains three main functionalities covering (1) implementations of published models, (2) benchmarking, and (3) data acquisition. A diagram of this is depicted in Figure 1, including the different utilities within each main functionality of *SUPR REF*.

**Figure 1:**
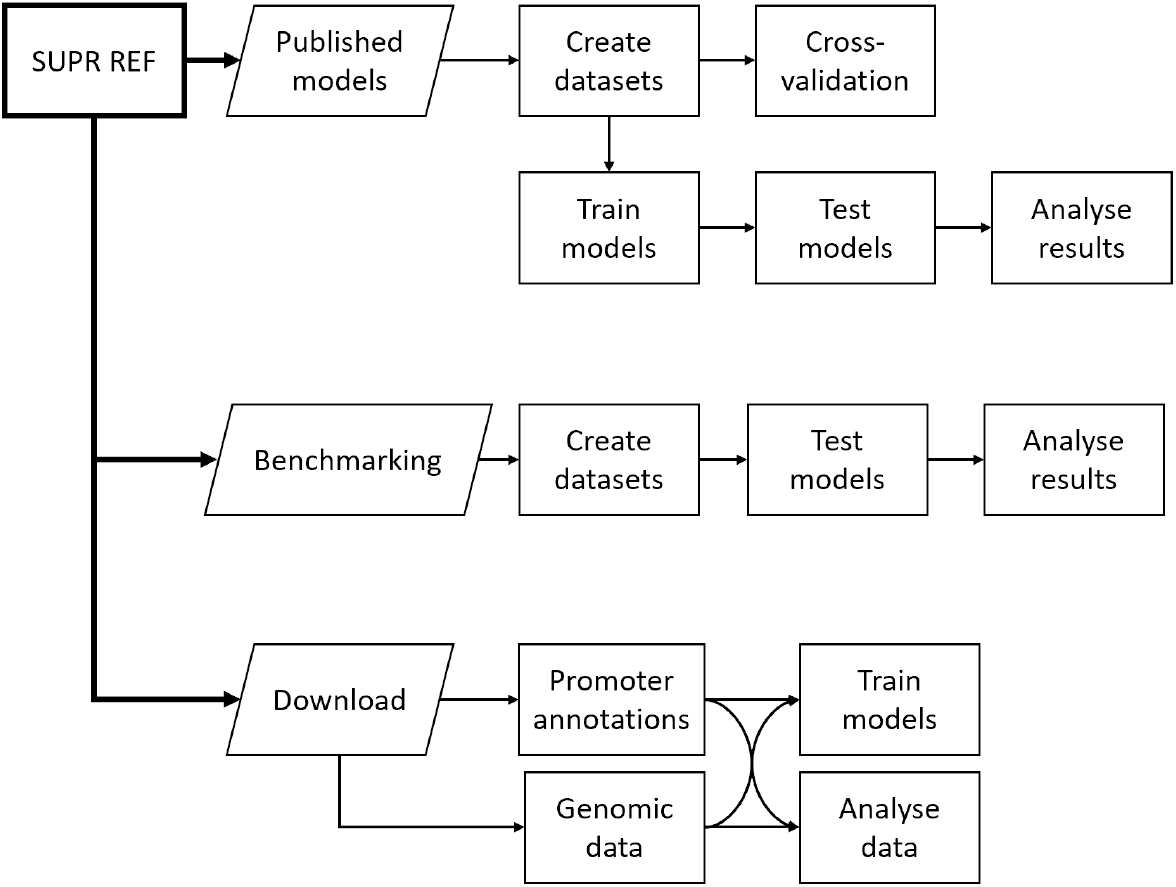
Schematic displaying main functionalities from SUPR REF

*SUPR REF* contains third party implemented architectures of recently published models, and the creation of their datasets. The datasets can be split into non-overlapping training and testing datasets between all models enabling proper model comparison. *SUPR REF* also includes training, testing, and cross-validation procedures for each model to analyse and visualise the models’ outputs.

The benchmarking system offers two main utilities, as well as analysis notebooks to help visualise and interpret how the trained models function. The `create` utility allows the creation of datasets from two approaches. The annotation approach utilises promoter annotation data and its corresponding reference genome to create a dataset. The sequence alignment approach creates a dataset by mapping the promoters in a genome using sequence alignment tools. Next, the `test` utility tests a trained DLPR model with previously published datasets or *SUPR REF*-created datasets. Two additional aiding utility are offered.

As a framework, SUPR REF can also aid in more general promoter recognition endeavours, including data acquisition and ML training with two additional utilities. The download utility is a data acquisition tool which aids in downloading promoter annotation data from biologically tested sources such as EPFL’s Eukaryotic Promoter Database by Dréos et al. [2017], UCSC’s upstream sequences by Haeussler et al. [2019] and Riken’s FANTOM 5 project by The FANTOM Consortium et al. [2014]. Finally, a `train` utility is available to help train promoter recognition models utilising DL approaches (user-made PyTorch modules) and other scikit-learn machine learning algorithms. These models can be trained on *SUPR REF*-created datasets which can be tailored to your specific needs within promoter recognition, such as promoter length and amount of upstream and downstream bases within the promoter region.

All utilities within *SUPR REF* include multiple parameters that can be adjusted based on the user’s needs, such as different dataset creation properties, and training and testing parameters. More information on each utility and its parameters can be found on the *SUPR REF* documentation.

### 2.2 DL models

As a recent field of study, DLPR models have not yet been systematically reviewed, although a recent overview can be found in the thesis by Perez Martell [2020]. In this work, we focus on the comparison of three recent DLPR models created for the human (hg38) genome. Human promoters are some of the most studied and annotated in the literature within eukaryotes. Therefore, human promoter annotations should contain adequately comprehensive data to successfully train an effective model.

The first method, **CNNProm**, was proposed by Umarov and Solovyev [2017] with a convolutional neural network (CNN) architecture. Promoter sequences were obtained from the Eukaryotic Promoter Database (EPD) by Dréos et al. [2017], while the non-promoter sequences were random subsequences within genes located after their first exons. Since the dataset is ambiguously described and only partially attainable, we obtained a dataset that to the best of our abilities resembled its description. The created dataset consists of promoter sequences from EPD that span 251 bases. Non-promoter sequences span the same length as promoters and are obtained from random locations after the first exon of each human gene’s sequence. To obtain these non-promoter sequences, we query the UCSC genome database by Haeussler et al. [2019]. Promoters were further separated into two classes and classified by different CNNProm models: promoters with a TATA box motif (TATA), and promoters without a TATA box motif (non-TATA). The CNNProm TATA model had a slightly different architecture from CNNProm non-TATA model. The main difference being the number of filters in the CNN and the pooling size, where TATA had a lower value in both cases. To reproduce their results, we used the previously created dataset with sequences of length 251. To properly compare them to other models, we used the same method of obtaining the sequences with an added 49 more bases upstream, totalling 300 bases for the sequences length.

Subsequently, **ICNNP** is a DLPR method by Qian et al. [2018], which was described as an improvement over CNNProm. Similar to CNNProm, ICNNP consists of a convolutional neural network architecture. Unlike CNNProm, which classified human sequences of length 251, ICNNP classifies sequences of length 300. This increase in length includes 49 additional bases in the upstream region of the promoter sequence. The ICNNP dataset includes a mix of TATA and non-TATA promoters from EPD, while the non-promoter sequences consist of human introns and coding sequences (CDS) collected from the 1998 GENIE dataset by Reese et al. [2000]. The non-promoter dataset of ICNNP was originally compiled by the Berkeley Drosophila Genome Project (BDGP) by The FlyBase Consortium [1999] for use as a representative benchmark dataset of human DNA sequences. To create ICNNP, the authors first focused on assessing the importance of various motif locations, referred to as ‘element sequences’, using *Support Vector Machines*. The predetermined locations were then extracted from the sequence to be used along a compressed (max-pooled) version of the complete sequence for classification by the neural network. Element sequences were found to account for most of the differentiative signal within promoters, while the compression of non-elements was found to help remove noise from the model. Therefore, the main difference between CNNProm and ICNNP was not from the neural network architecture, but from the pre-processing of the sequences.

The final DLPR method, **DeePromoter (DProm)** (Oubounyt et al. [2019]) consists of a convolutional long short-term memory (CLSTM) architecture. Similar to work by Umarov and Solovyev [2017], Oubounyt et al. created different models for promoters with and without a TATA box motif, along with model separation by species. Promoter sequences were obtained from EPD; for non-promoter sequences, DProm used synthetically created sequences by changing the promoter sequences. DProm models classify human sequences with a length of 300 bases.

All models encode DNA sequences using one-hot encoding, i.e., a matrix where each of the rows consists of unique bases (A, C, G, T) and the columns are the locations in the sequence. The only difference is found in ICNNP where the sequences are further separated into ‘element’ and ‘non-element’ subsequences before one-hot encoding.

Each of the previous DLPR methods was assessed on a different dataset. The dataset for CNNProm contained 57,224 DNA sequences, comprising of 9,682 sequences (14.72% promoter) for ‘Human TATA’ model and 47,542 sequences (41.67% promoter) for ‘Human non-TATA’ model. The dataset for ICNNP contained the least amount of sequences with 12,391 sequences (57.75% promoter). Finally, the dataset for DProm contained 59,194 sequences comprising of 6,130 sequences (50% promoters) for ‘Human TATA’ model and 53,064 sequences (50% promoters) for ‘Human non-TATA’ model. The specific number of promoters and non-promoters for all three methods is available in Table 1.

**Table 1:**
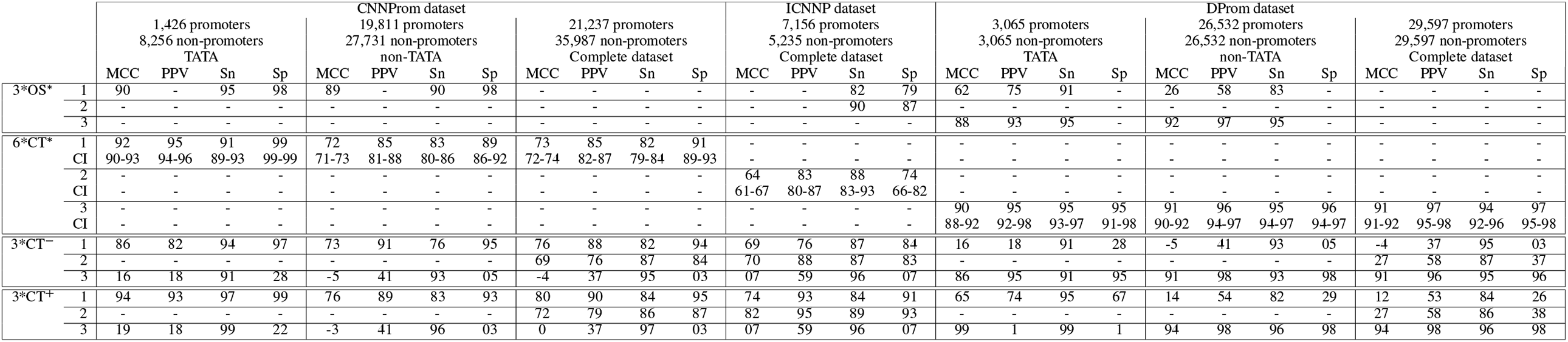
Performance results from human DLPR models. OS indicates model results from original studies. CT shows cross testing results from our implemented models. (1) CNNProm, (2) ICNNP, (3) DProm model. CT^+^: results from training and testing datasets with some overlapping sequences. CT^−^: results from properly split training and testing datasets having no overlapping sequences. CT^∗^ and OS^∗^: results obtained from 10-fold cross-validation. All values expressed as percentages. 99% confidence interval values for our 10-fold cross validation results shown within CI rows.

### 2.3 Benchmark Datasets

*SUPR REF* contains two dataset creation approaches. The sequence alignment approach receives as input promoter sequences and requires a local alignment tool (here: BLAST) to align them to a genome. When exact promoter sequences (without mutations or indels) are found in a genome, this approach might not be as accurate or efficient as exact string matching algorithms. Therefore, we recommend using this approach when annotations are unavailable, or when there are slight variations in the genome or promoter sequences. The annotation approach requires annotations containing the TSS locations of genes. The annotations for this study were obtained from MGA database by Dréos et al. [2018]. It obtains its data from multiple sources including EPD. The exact genome annotation used is called `Hs_EPDnew_006_hg38`^1^. With the TSS locations denoted as +1, a promoter region is extrapolated by obtaining 249 bases upstream and 50 bases downstream of the TSS. Therefore, the promoters span regions from −249 to +50 for each TSS in the annotations. The annotation approach was chosen to create the benchmark datasets in this study, as the data was readily available for the hg38 genome.

With *SUPR REF*, we created multiple benchmark datasets using the annotation approach and a sliding window over the hg38 genome. To resemble the datasets for previous models, our dataset creation parameters follow a stride of 50 and a window size of 300. Any window with an ambiguous base ‘N’ was removed to avoid model noise and bias. Sequences were categorised as promoter if at least 250 out of 300 bases (83%) were contiguously overlapping with true promoter sequences. All other sequences were categorised as non-promoter. These benchmark datasets follow the IO tagging scheme, where “I” means inside (promoter sequence) and “O” means outside (non-promoter sequence). Tagging schemes stem from the Natural Language Processing (NLP) community (Ju et al. [2019]). They translate well into promoter recognition where DNA is regarded as a text corpus, and promoters as entities within that corpus.

To elicit a more realistic performance of the DLPR models, we created a larger testing dataset comprising of hg38 chromosome 1 and 2 (hg38chr1&2). This larger dataset contains 422,912 sequences, as opposed to less than 60,000 sequences from each study’s cross-validation test. The dataset contains overlapping sequences designed to test sequences where a few bases on the sequences’ edges are different. This could alert us if a model has strict positional learned behaviour or if it is overfitting on specific sequence patterns. The hg38chr1&2 dataset was split into its respective chromosomes. The hg38 chromosome 1 (hg38chr1) benchmark dataset includes 196,224 DNA sequences with 3,748 (1.91%) promoter sequences. These sequences consist of multiple known promoters from EPD for the approximately 2,000 known coding genes in the human chromosome 1, and also the overlapping sequences that occur from the sliding window approach. The hg38 chromosome 2 (hg38chr2) benchmark dataset includes 226,688 DNA sequences with 2,632 (1.16%) promoter sequences. Notably, it is larger than hg38chr1 even though chromosome 1 contains more base pairs. This is because of the higher predominance of ‘N’ bases in chromosome 1 annotation. The sequences for hg38chr2 also consist of multiple known promoters from EPD for the approximately 1,200 known coding genes in the human chromosome 2, and also the overlapping sequences that occur from the sliding window approach.

A fair testing procedure is incomplete without a held-out testing dataset that is completely unseen by the trained model. Since some models are being trained on the complete set of human promoters, it is important to test them on a dataset where human promoters are not available to ensure no overlapping occurs within the training and testing datasets. Therefore we also include a dataset of mm10 chromosomes. As with the human benchmark dataset, the mm10 dataset contains sequences from chromosome 1 and 2 of mice. The mm10 chromosome 1 (mm10chr1) benchmark dataset includes 191,093 DNA sequences with 3,275 (1.71%) promoter sequences, while the mm10 chromosome 2 (mm10chr2) benchmark dataset includes 193,183 DNA sequences with 4,129 (2.13%) promoter sequences.

### 2.4 Model implementations

Our implementation of CNNProm follows the work by Umarov and Solovyev [2017]. To reproduce the published results, we trained our implemented CNNProm model utilising sequences with a length of 251 bases. Afterwards, to properly compare our implemented CNNProm model to other DLPR models, we used sequences with 300 bases to match the other two models which were originally designed for use with 300 bases. To create a model that encapsulates both TATA and non-TATA promoters, we decided to use the non-TATA architecture, which contains the highest amount of CNN filters between the two models. In the context of promoter recognition, CNN layers are used to find motifs within the sequence regardless of their locations within the sequence. With this architecture, we implemented the CNNProm model for this study using the same configuration as described by Umarov and Solovyev [2017] and shown in Figure 2a.. In *SUPR REF*, a CNN layer for motif discovery comprises of a user-specified number of filters of length 15 and 21 for finding differently sized motifs, followed by a max-pooling operation and a rectified linear unit (ReLU) activation function. This common CNN architecture can be expanded by increasing the number of CNN layers on top of each other. Additionally, each CNN layer can make use of normalization and dropout.

**Figure 2:**
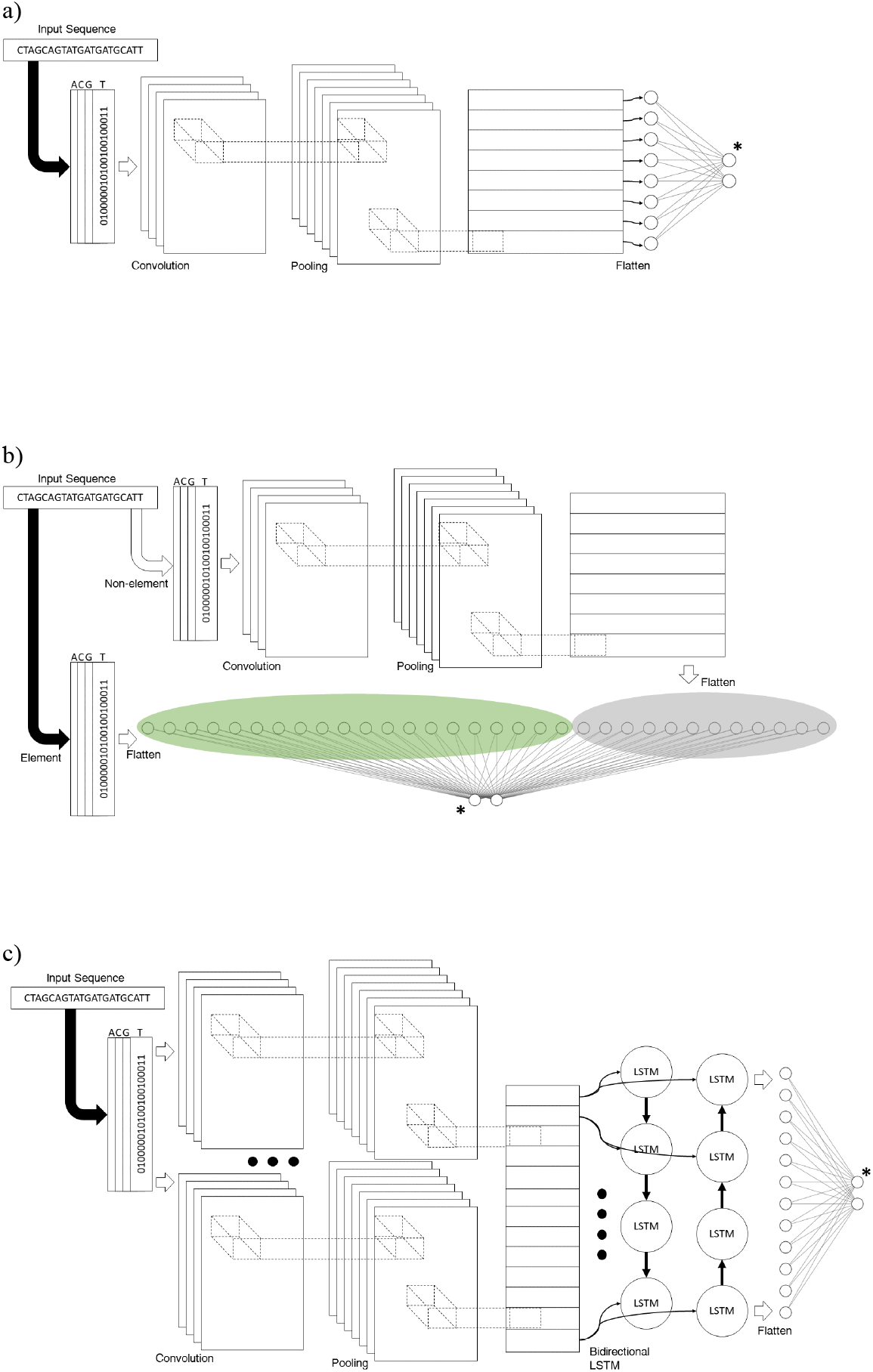
Model architectures from a) CNNProm, Umarov and Solovyev [2017], b) ICNNP, Qian et al. [2018], c) DProm, Oubounyt et al. [2019]. ^∗^Two neurons depicted as output layer for softmax activation, but a single output neuron is used in case of sigmoid activation.

Our implementation of ICNNP follows the work by Qian et al. [2018]. The input sequences contain 300 nucleotides, and the architecture uses a similar CNN layer to CNNProm containing 200 filters and a max-pool layer, with the addition of a parallel layer containing location-specific ‘element’ subsequences that are concatenated to the CNN layer’s output to aid in the classification process. This architecture is shown in Figure 2b. This architecture is clearly dependent on nucleotide positions within the 300 length sequence, which at a genome level will require a sliding window approach to use a very short stride. This means that the model will have to be run once for almost every base in a genome, which in the human genome would amount to more than 6 billion times, making it infeasible for use in a genome wide promoter recognition setting.

Our implementation of DProm follows the work by Oubounyt et al. [2019]. The input sequences contain 300 bases, and this architecture utilises a combination of CNN and recurrent neural network (RNN) layers. This architecture is shown in Figure 2c. The CNN layer acts as an embedding layer to locate the motifs that get passed to a bidirectional long short-term memory (LSTM) RNN layer that interprets the sequential signal of the data, namely the order in which motifs are located within the sequence.

#### 2.4.1 Data cleaning

While implementing the models and recreating their datasets, we noticed that some promoters from EPD appeared multiple times within the same dataset. This could lead to biases in the training and testing datasets. To avoid this, we removed any duplicate sequence after the first occurrence. For example, the CNNProm non-TATA dataset contains 687 duplicate sequences, which can translate to ∼ 2 − 5% difference in MCC values.

We further created a script to completely separate all testing and training datasets for all three models for our cross testing evaluation. To do this, we first generated separate training and testing datasets for DProm since it contains all promoter sequences. Afterwards, the training and testing datasets are further reduced to match the number of promoters needed by CNNProm and ICNNP. The non-promoter sequences are then appended to the training and testing datasets to match the complete amount of data needed to train each model. Finally, we make sure that no sequences are overlapping between training and testing datasets for all three models by comparing the rows from each dataset.

### 2.5 Model benchmarking

Our training and testing procedures were implemented using Skorch (Tietz et al. [2017]), a library adding functionality to PyTorch. The most useful included features are the ability to use training, testing and cross-validation, and evaluation metrics supported by Scikit-learn (Pedregosa et al. [2011], Buitinck et al. [2013]). This makes *SUPR REF*’s implementation easy to maintain, and training and testing procedures consistent.

The training process involves importing a *SUPR REF* created dataset and using stratified sampling to create the training batches. The models can be trained with two different output functions: sigmoid and softmax. Sigmoid outputs a single value that can separate promoters and non-promoters by a threshold. Softmax outputs two values: the probability that the input sequence is or is not a promoter. The loss function follows the output function: sigmoid utilises binary cross entropy loss. Softmax utilises cross entropy loss. Model training is completed once one of the following two conditions is met: 50 epochs have occurred or the performance of the model has not increased in 5 epochs. Each epoch produces a checkpoint file with the best performing model at that time. The checkpoint file contains all the model’s parameters, or each neuron’s values in the neural network architecture. The learning rate for all models was set to 0.001. Batch size and optimizer differed depending on the original model’s description.

The testing process imports a dataset and loads the best performing checkpoint of a trained model. Next, the trained model is used to predict whether the sequences from the imported dataset are promoters or non-promoters. The predictions are then compared to their sequences’ true classification, and evaluation metrics are calculated.

### 2.6 Evaluation metrics

Choosing the proper metrics to evaluate supervised machine learning models is a crucial step for testing the reliability to perform the task that the model has been trained to do. Metrics also help to compare different trained models when tested on the same dataset. In the case of promoter recognition, the problem can be represented analogously to the classification of DNA sequences. Therefore, promoter recognition can be designed as a binary classification task that can be evaluated—such as in related work by Oubounyt et al. [2019], Qian et al. [2018], Umarov and Solovyev [2017]—with the following metrics: Sensitivity (Sn), Specificity (Sp), Precision (PPV), and Matthews correlation coefficient (MCC).

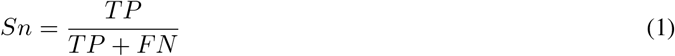

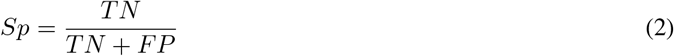

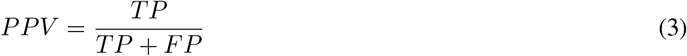

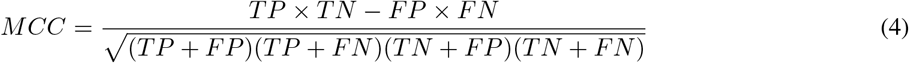

Here: *TP* is the number of true positives (i.e., correctly identified promoter sequences); *TN*, true negative (correctly rejected promoter sequences); *FP*, false positive (incorrectly identified promoter sequences); *FN*, false negative (incorrectly rejected promoter sequences).

## 3 Results and discussion

Here we provide results from three DLPR models and their comparisons to our own implementations. These are followed by testing our implementations on larger benchmark datasets that more closely resemble the imbalanced proportion of promoters in nature. The benchmarking procedure demonstrates a lack of precision in the models, leading us to investigate common tactics to mitigate it. These tactics include the use of various synthetic data to augment the training dataset, exploration of different output functions in neural networks for classification of promoter sequences, and sampling methodologies on training data. We include results on the impact different promoter annotation datasets can have on a model’s ability to recognise promoters, and provide an analysis of previously utilised datasets for DLPR and highlight how a model’s performance metrics can entirely depend on a small portion of the dataset. We then present the analysis toolkit for trained models, which can analyse the weights in CNN and RNN layers.

### 3.1 Initial comparisons

Looking at the datasets used to train the published models, we notice that non-promoter sequences are the main differentiating factors affecting the training process. To investigate the extent that the different negative sequences affects these datasets, we test each of the previous DLPR models on human datasets from other studies. We refer to this as *cross testing* the models. First, we mixed the complete human dataset—including TATA and non-TATA—of each study to create a complete training dataset for each corresponding model. In essence, we implemented and trained each model on its study’s set of available datasets containing sequences of length 300, including a complete dataset merging TATA and non-TATA promoters. Then, each model was tested on all the datasets for a proper comparison of the models.

Results are provided in Table 1, showing performance metrics of each model when tested on their own dataset, alongside our implemented cross validation results. Each model’s results from a published study is contained within the **original study** (OS) row. Our implementation results are found under **cross testing** (CT) rows. CT rows include models trained on sequences that overlap (*CT* ^+^) with sequences on the testing dataset and models trained on proper train-test split (*CT* ^−^) datasets; this shows the effect that improper splitting can have when evaluating models. For easier visibility, CNNProm, ICNNP, and DProm have been renamed to 1, 2, and 3 respectively. From Table 1, we see that OS (1) CNNProm has been tested on all three datasets, while OS (2) ICNNP and OS (3) DProm were tested only on their own dataset. This is especially problematic as OS (2) ICNNP and OS (3) DProm compare their cross-validation results with OS (1) CNNProm’s results, but it is unclear how OS (1) CNNProm was trained or tested on the other datasets. This unknown procedure makes it difficult to assess the fairness and validity of the comparison between the models.

Results from our cross validation (*CT* ^∗^) did not exactly match the OS results. This is specially evident in the CNNProm non-TATA and complete models, as CNNProm was originally designed for sequences of length 250 and the architecture is highly tuned for that length. We observe that some OS results resemble *CT* ^+^ more closely than those from properly split datasets i.e., ICNNP model being superior to CNNProm model on the ICNNP dataset. Notably, the CNNProm non-TATA model did not obtain the same performance as in the original publication, even with overlapping and duplicate sequences when trained on sequences of length 300, which prompted us to examine the original CNNProm non-TATA dataset further. We found that the original dataset includes 686 duplicate sequences out of 47,541 sequences that can skew cross-validation, while our implemented dataset creation for CNNProm did not include duplicate sequences.

When comparing results from OS CNNProm tested on CNNProm dataset and ICNNP dataset, we observe a discrepancy for the CNNProm model: CNNProm performs better on the CNNProm complete dataset, while ICNNP outperforms CNNProm on their ICNNP complete dataset. The inferior performance of CNNProm seems to be occurring because of the improperly split training and testing datasets, as the properly split dataset has very similar performance to ICNNP on MCC. This discrepancy in performance could also be due to the difference in non-promoter sequences since the promoter sequences for ICNNP dataset are a subset of the CNNProm dataset promoter sequences. Similarly, results from OS CNNProm on the DProm dataset differ significantly from its performance on the CNNProm dataset. As previously mentioned with the CNNProm and ICNNP datasets, promoter sequences are very similar in both CNNProm and DProm datasets; in this case, CNNProm promoter sequences being a subset of DProm promoter sequences. Therefore, the difference in CNNProm’s performance on the DProm dataset can be mostly attributed to non-promoter sequences.

The conflicting performance results from all three models when tested on datasets from different studies show that cross-validation results on small datasets are insufficient to capture a model’s true performance. We infer that discrepancies in test results might stem from a limited number of training samples. This can lead to insufficient patterns to adequately represent the distribution of promoter and non-promoter sequences within real DNA sequences found in nature. As such, a more representative dataset should be used to validate the true performance of these models.

### 3.2 Benchmark comparison

We were able to successfully reproduce the results for all three models by obtaining highly similar cross-validation performance on their respective datasets. Implementation results to verify the validity of our implemented models can be obtained from our framework, which is available online at GitHub. All three implemented models were trained and tested on the a common promoter database (EPD), meaning that these promoter sequences appeared in both the training and testing datasets of each other without proper consideration. This bias will skew the sensitivity and precision metrics towards more positive values. Some non-promoter sequences in certain training sets (CNNProm, ICNNP) are also found in the testing datasets, which will skew specificity and MCC in the same manner as previously stated. Therefore, all trained models were also tested on hg38 benchmark dataset. The two most recent models were also tested on the smaller hg38chr1&2 and mm10chr1&2 benchmark datasets. The results of the models being tested on the larger, more representative human genome dataset and the smaller human and mouse chromosome datasets are shown in Table 2. Note that the precision of all three models decrease to nearly zero in the hg38 dataset. This implies that many non-promoter sequences are being identified as promoters by these models, which highlights the limited capability of the models when dealing with larger sets of data that resembles the promoter to non-promoter ratio in nature. It is also important to notice that models are able to perform similarly within related species, as the models perform comparably in both human and mouse chromosomes. Finally, note how precision generally gets lower as the amount of non-promoter sequences increases, which occurs from the increasing number of false positives.

**Table 2:**
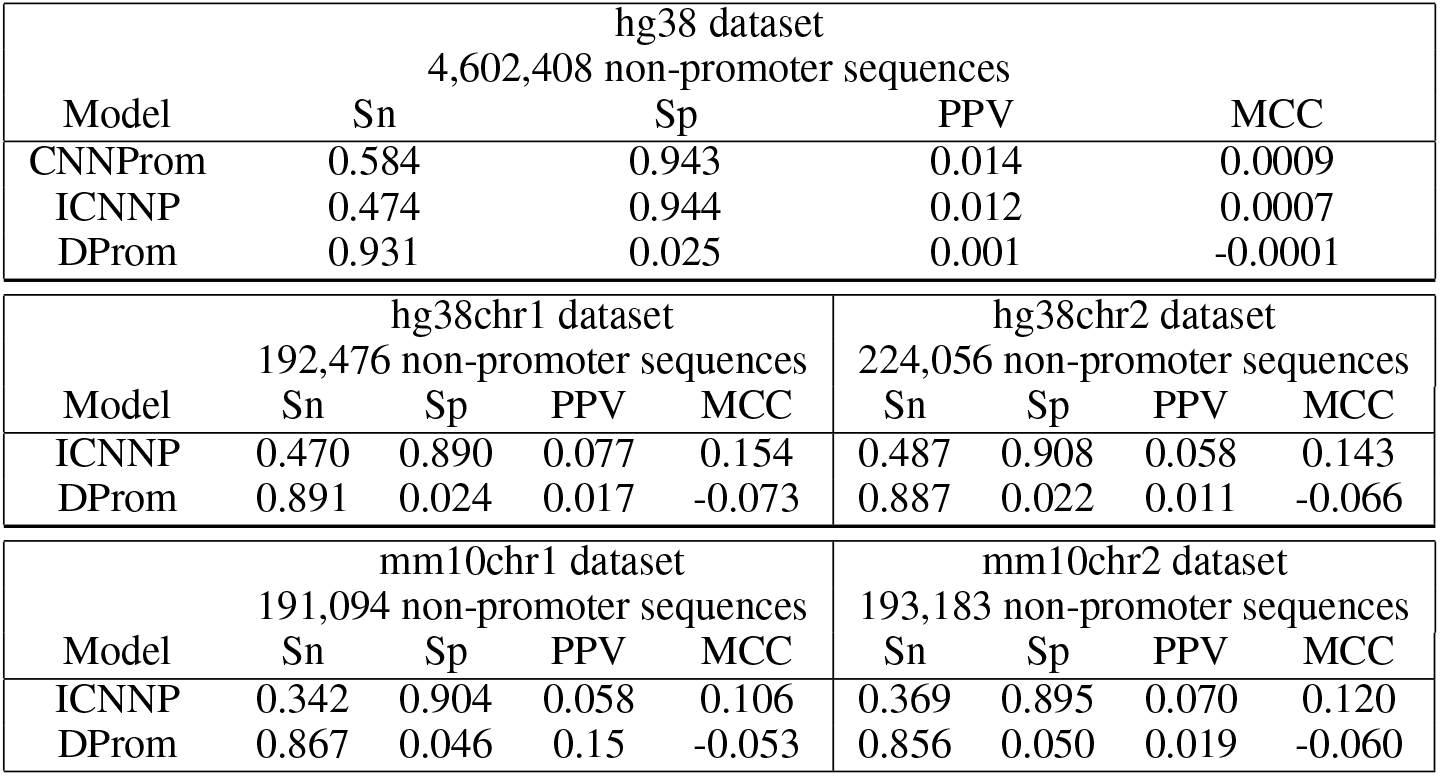
Results from implemented models when tested on multiple benchmark datasets. Models are clearly lacking in precision (PPV).

### 3.3 Synthetic data

We further explored promoter modification into non-promoter sequences by observing the effects of the modified base distribution on the training of models. The models had DProm architecture and were trained on multiple different synthetic non-promoter datasets. In the DProm study, substitutions occurred in a uniform manner, i.e., any base had the same probability of being randomly substituted. Here, we calculated the original promoters’ base distributions and made the substitutions match the original promoter distributions, i.e., the bases were shuffled instead of replaced by independent DNA subsequences. Therefore, the two types of synthetic data tested are the ‘uniform distribution’ (UD) and ‘original promoter distribution’ (OPD), respectively. When training only on TATA promoters, the models had difficulty differentiating the original promoter distribution sequences from non-promoters, leading to a lower sensitivity compared to the uniform distribution sequences. The sensitivity increased for non-TATA promoters, although uniform distribution models still outperformed the original promoter distribution. The lower sensitivity was traded for slightly higher specificity, although MCC remained generally slightly lower for OPD than UD trained models. These results can be seen in Figure 3a. These experiments validate the use of uniformly random distribution substitutions rather than original promoter distribution substitutions as done by Oubounyt et al. [2019].

**Figure 3:**
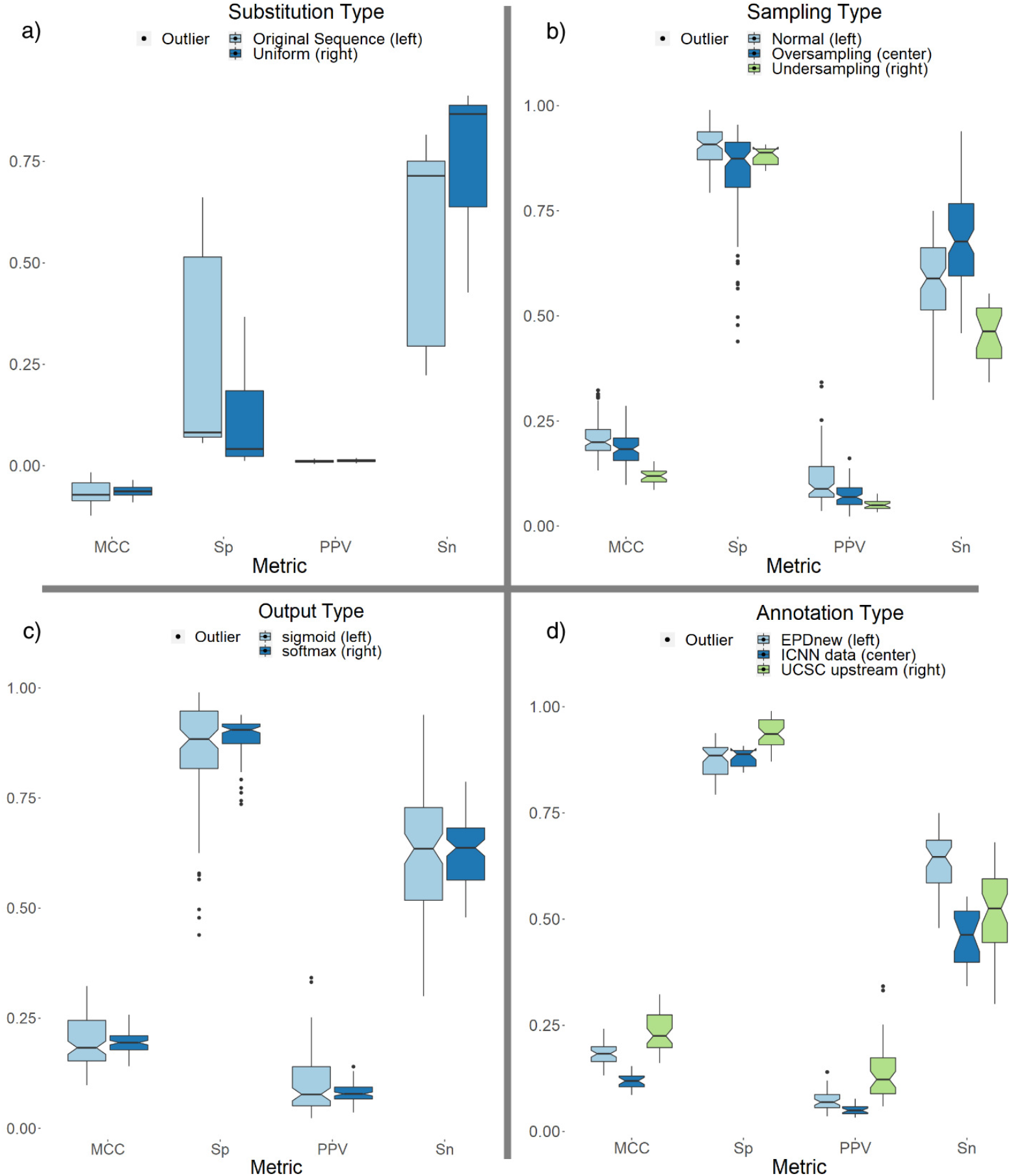
Results from experiments in a) Synthetic data, b) Sampling methods, c) Output functions, d) Annotations

While non-promoter sequences created from OPD helped models obtain slightly higher specificity than UD sequences when tested on hg38chr1, using non-promoter synthetic data to train models had disadvantages over non-synthetic data: all models trained on synthetic data had lower specificity than models trained on non-synthetic data.

### 3.4 Sampling methods

Common methods to compensate for highly imbalanced datasets when training models include oversampling and under-sampling. Oversampling is generally done by duplicating the minority class within the training dataset, undersampling removes samples from the majority class. Both of these methods try bringing the number of promoter and non-promoter sequences to a similar value to aid in model training. We trained a model, referred to as ‘normal sampling’, with the benchmark hg38chr1 which contains around 100 non-promoters to each promoter sequence in the dataset. We then trained a second model referred to as ‘undersampling’. This model contains at most 10 non-promoter sequences for each promoter in the dataset by limiting the number of non-promoter sequences. Finally, we trained a third model referred to as ‘oversampling’, trained with a ratio of one promoter per 10 non-promoter sequences—in this case the promoter sequences were duplicated to achieve that ratio. Comparing the results from these three sampling methods, shown in Figure 3b, we observe that the normal sampling model outperforms the other two methods in MCC, precision and specificity. The oversampling method had higher sensitivity than the other sampling methods.

### 3.5 Output functions

We explored how the activation function in the output layer of a neural network can affect the performance of models. The most commonly used output activation functions are sigmoid and softmax. When using sigmoid, a numerical threshold is required to separate the classification between promoters and non-promoters. The threshold in our experiments for models utilising a sigmoid activation function is 0.5. This threshold is unnecessary for a softmax function since it outputs the probability of a sequence being a promoter or non-promoter. Results for this experiment is shown in Figure 3c. Notably, the sigmoid function in most of our models tends to provide a small performance increase in precision while the softmax function tends to provide increased sensitivity, and less fluctuation within all metrics.

### 3.6 Annotations

Promoter annotation datasets contain promoter sequences that have been biologically verified. The EPDnew database has been utilised in most promoter recognition studies. We compare these annotations to UCSC’s upstream datasets, which contain upstream sequences for every biologically verified gene in multiple organisms. These upstream sequences can be used to obtain promoter sequences since they are adjacent to the TSS of their respective genes. It is important to note that the upstream dataset cannot account for alternative TSS of genes that occur frequently in nature. Results for this experiment are shown in Figure 3d. When tested on benchmark datasets, models trained on UCSC upstream data had higher precision than models trained on EPDnew promoters. The trade-off to the higher precision in models trained on UCSC upstream data was of lower sensitivity.

### 3.7 Analysis

To understand how the models could differentiate between promoter and non-promoter classes, *SUPR REF* allows us to analyse the sequences of the datasets. This helps us choose the relevant positions of the sequence to extract. These subsequences can then be used to train and test models. In the non-TATA dataset used for training CNNProm models as seen in Figure 4a, promoter sequences tend to contain minor differentiating motifs in the regions from –22 to −10, and +17 to +31, where +1 is the TSS located at position 201 of the sequence. A more pronounced differentiating region can be clearly located from −4 to +2. Looking at the TATA dataset in Figure 4b, we find that the TATA box motif to be a clear differentiating factor ranging from –35 to −20. We can also find minor differentiating bases between –200 and −40. We explored whether a smaller stretch of DNA would be enough to create a model that performs similarly to the results of the CNNProm study by training a model with only the −35 to +2 subsequence. Note that this includes a significant differentiating region from −2 to +1. We found that this limited model performs similarly to models utilising the complete sequence. To validate the impact that training and testing datasets can have in the evaluation of a model’s performance, we further explored using only the 10 bases spanning the TATA box motif to train a model (10-bases model). The TATA box is found in the regions between –33 to −23, and using only these 10 bases in a cross-validation experiment, we achieved the results shown in Table 3, which are comparable to the results from OS CNNProm in Table 1. This shows us that minor differentiating regions such as –200 to −40 and very short but significant differentiating regions such as −2 to +1, did not affect the performance of this model because a much longer and highly significant region was still present within the training dataset.

**Figure 4:**
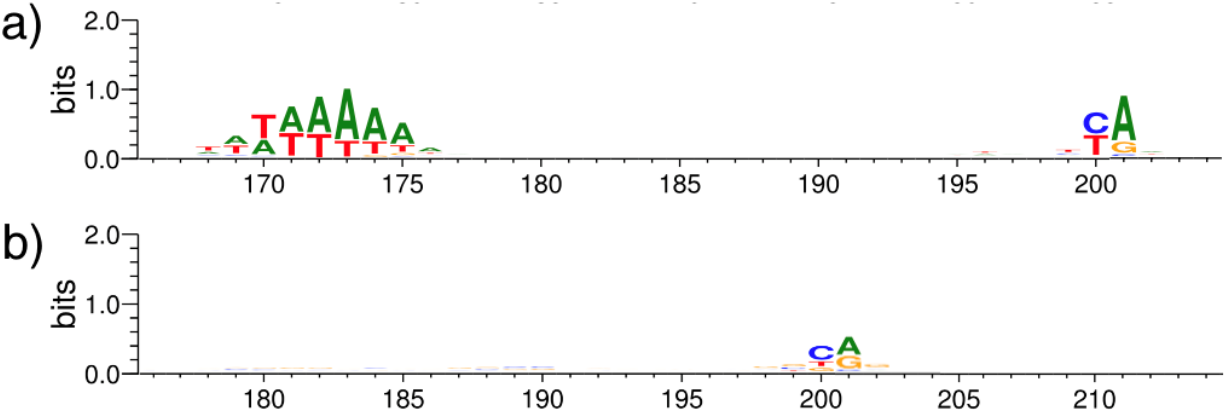
Differentiating bases from promoters in a) CNNProm TATA dataset, b) CNNProm non-TATA dataset. Notice: tiny letters are included within the seqlogos, e.g., position 1 to 40 in (a) and 179 to 184 in (b). Clear TATA box motif located near position 170 to 175 in (a).

**Table 3:** CNNProm model trained and tested on 10 bases of the human TATA dataset. Metrics comparable to Table 1 OS CNNProm TATA dataset results.

| Model | Sn | Sp | PPV | MCC |
| --- | --- | --- | --- | --- |
| CNNProm 10-bases | 0.95 | 0.95 | 0.91 | 0.89 |

### 3.8 Analysis toolkit for trained models

Models from the studies examined here categorise sequences into being promoters or non-promoters. The output from these models do not provide a way to further explore what features the models might be using to make this decision. Therefore, we created an analysis toolkit for CNN and RNN layers within trained models that can highlight the bases and locations that carry the most weight into the final output. We also created a model that can import JASPAR motifs as CNN filters to analyse where they appear in promoters and non-promoters. This model was trained with the CNNProm non-TATA dataset. The results from the analysis give multiple seqlogo visuals for each filter (JASPAR PolII motif), as well as visuals for the complete sequence; a combination of all the filters in the sequence provides a global view of the bases having an impact in the final output. Figure 5 shows an example of an imported JASPAR motif, specifically the human *initator element* (INR). Notice that the filter weights seem very small because of normalisation within the neural network, which causes some bases to not appear in the seqlogo in comparison to the JASPAR seqlogo^2^. Figure 6 depicts the location where the model found INR within promoter sequences in CNNProm human non-TATA dataset. Lastly, Figure 7 shows which bases the model is looking at using all human JASPAR motif filters combined.

**Figure 5:**
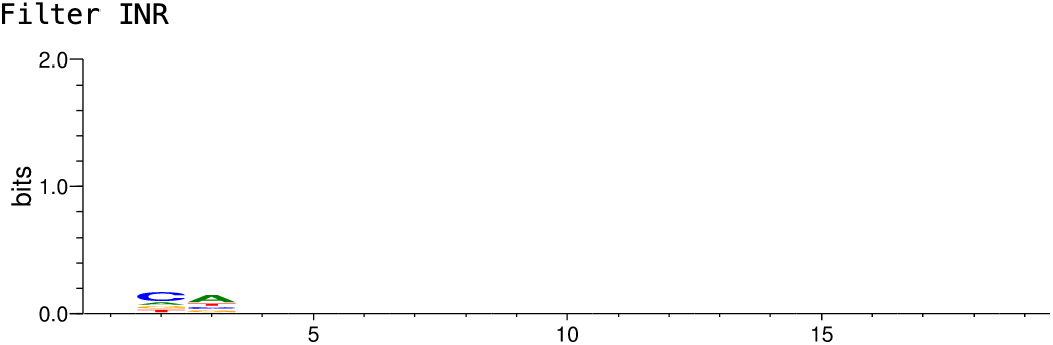
Seqlogo showing the filter weights in a CNN layer for the INR motif imported from JASPAR.

**Figure 6:**
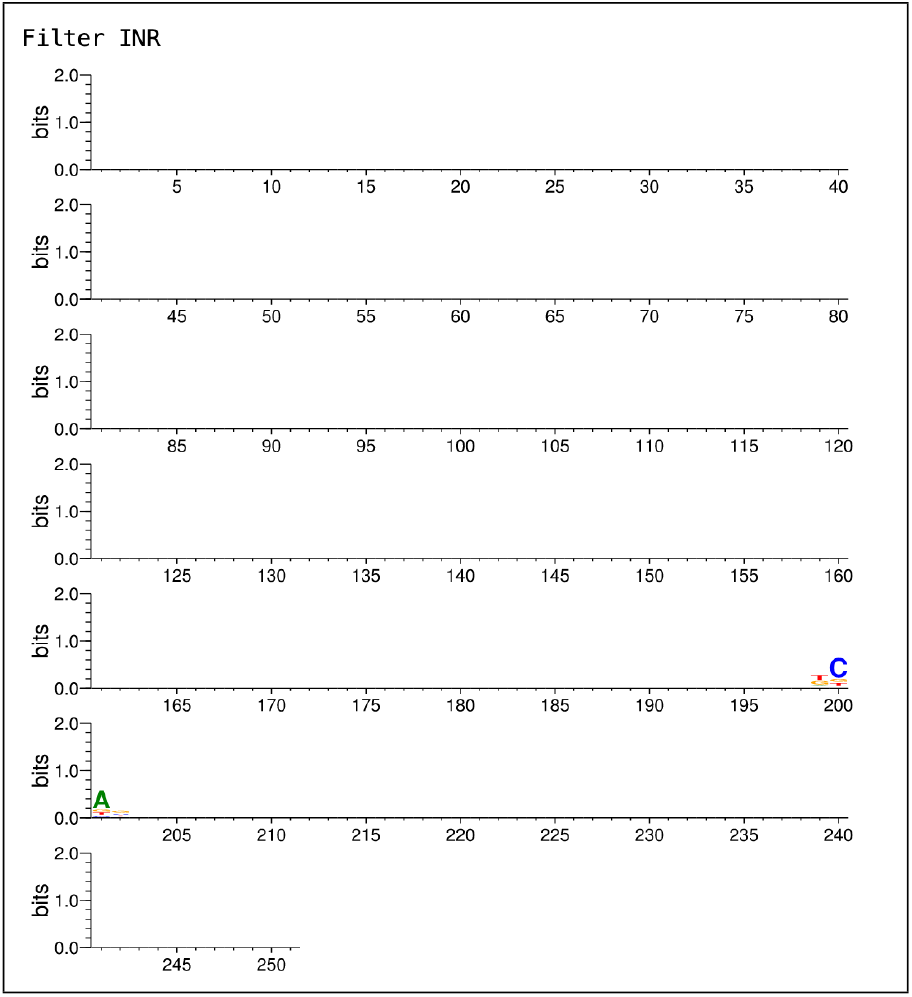
Seqlogo showing where the model is locating the INR motif within the promoter sequences of CNNProm human non-TATA dataset.

**Figure 7:**
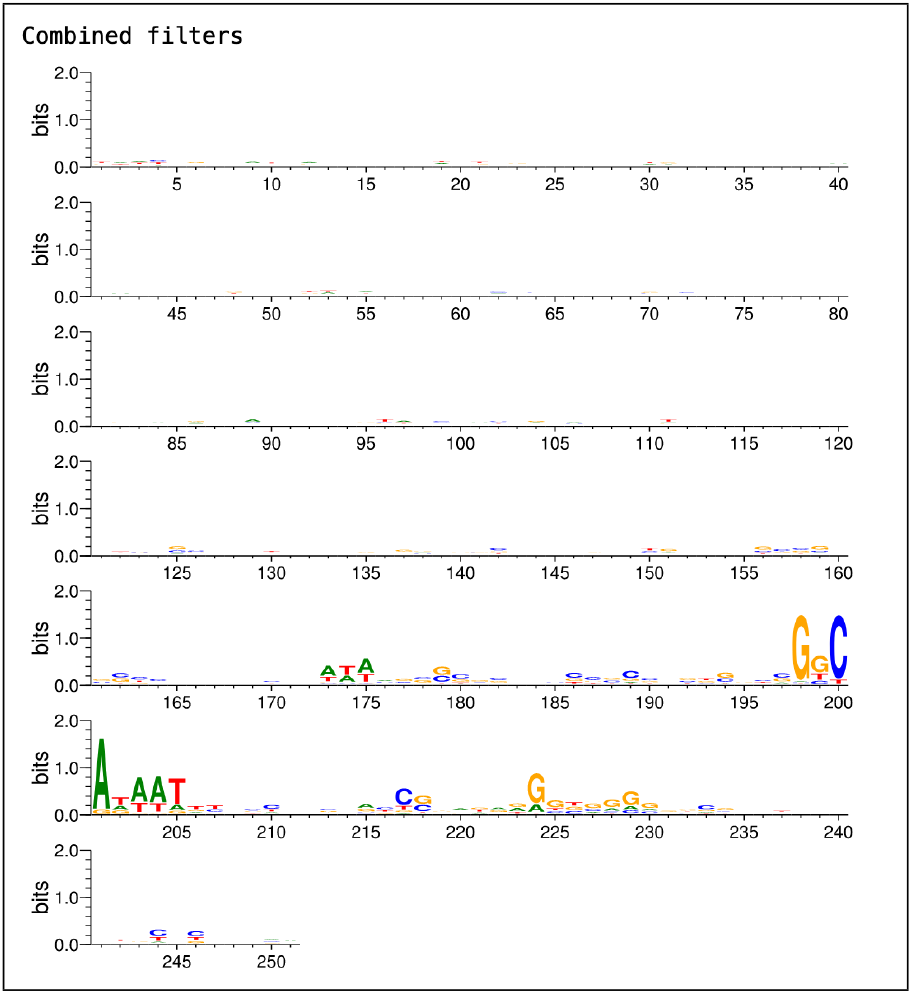
Seqlogo showing the bases that are preferred by all the CNN JASPAR-based filters combined within promoter sequences of CNNProm human non-TATA dataset.

## 4 Future work and conclusions

We presented that the proper assessment of promoter recognition models can be heavily impacted by both the training and the testing datasets that are assumed to be a sufficient sample size of the DNA sequences observed in nature. Although the three studies we examined utilised a similar set of promoter sequences from EPDnew, they did not use the same dataset for a proper comparison of their models. Selecting the best promoter recognition model can therefore be difficult to assess using the results from the published studies or, worse still, can lead to the wrong conclusion if choosing an under-performing model. To tackle this problem and bring clearer comparisons to assess DLPR models, we created *SUPR REF* and included the three literature model architectures, training procedures, as well as tools for benchmark dataset creation and their subsequent analysis.

We showcase *SUPR REF* by comparing previously published DLPR models in a highly imbalanced dataset (resembling ratios found at the genomic scale), alongside several experiments to explore different training methods that can increase performance. We provide a more specific promoter recognition analysis and data that help carry and accelerate DLPR research. Our results indicate that the true performance of *ab initio* promoter recognition models is still not at an acceptable level for highly imbalanced datasets, as precision is severely lacking. Nevertheless, our framework can shed light on the underlying methods of a model that recognises promoters within highly imbalanced data and between related species.

### SUPR REF

is a framework that aims to be at the forefront of reproducible research within deep learning for bioinformatics. Within the last decade, an ongoing debate within the scientific community regarding reproducibility problems has come to light (Fidler and Wilcox [2021]). This debate has also permeated computational fields, as shown by Hutson [2018], where data should be more accessible and distributed. The bioinformatics field is no exception to this problem, and current technology can help mitigate this (Kim et al. [2018]).

Our framework makes use of Scikit-learn, which simplifies the inclusion of state-of-the-art algorithms for training and testing models. Further improvements for benchmarking DLPR models can be achieved from other recently developed testing techniques, such as nested cross-validation by Bates et al. [2021], made available by Scikit-learn and other similar libraries which *SUPR REF* used for its development.

Following our results, we hypothesise that training models with a mix of promoters from multiple related species could potentially be a mitigation strategy for the imbalance in training datasets of promoter recognition within that branch of life. This works in accordance to how models trained on mouse promoters can also perform similarly on human promoters without the need of retraining the model. The inclusion of more promoter sequences can also increase performance as found with the UCSC upstream annotations containing 60,555 sequences as opposed to approximately 30,000 from EPD. These ideas can be further combined by training a model on different annotations and multiple related species.

Increasing the dataset size of promoter sequences could be possible using non-reference genomes. Once the myriad of biologically functioning variants of promoter sequences is identified, they can be clustered together by similarity. This can help the training process of DLPR models differentiate between the distribution of base pairs that are involved in promoters sequences and non-promoter sequences, thus avoiding noise that might occur in the case of IO tagging scheme. As such, promoters would not only be categorised as a single class, but as multiple classes of promoters depending on the clustering. Techniques to further improve performance of DLPR models are incrementally developed from best performing ideas and designs. It is imperative that frameworks like *SUPR REF* simplify the process of improving models by removing the need to implement previously known techniques and being hindered by implementation details. This work is a step toward this goal while we increase our understanding of promoter sequences necessary for the transcription process that occurs within cells.

## Footnotes

1 https://ccg.epfl.ch/mga/hg38/epd/Hs_EPDnew_006_hg38.sga.gz

2 http://jaspar.genereg.net/matrix/POL002.1/

